# The bacterial schizorhodopsins: novel light-driven inward proton pumps from Antarctic *Minisyncoccota* (Patescibacteria) and cyanobacteria, with implications for the proton-pumping mechanism

**DOI:** 10.1101/2025.10.15.682728

**Authors:** María del Carmen Marín, Masae Konno, Andrey Rozenberg, Oded Béjà, Keiichi Inoue

## Abstract

Microbial rhodopsins represent a diverse superfamily of light-sensitive seven-transmembrane proteins with expanding phylogenetic diversity driven by advances in metagenomics. Among these, schizorhodopsins constitute a divergent family originally identified as inward proton pumps from *Promethearchaeota* (Asgard archaea). Here, we report that in addition to archaeal schizorhodopsins, many members of the family originate from bacteria and detail a comprehensive biophysical characterization of novel schizorhodopsins from Antarctic *Minisyncoccota* (Patescibacteria) and cyanobacteria, designated as paSzR and psSzR, respectively. Both proteins function as light-driven inward proton pumps, as confirmed through pH measurements in *Escherichia coli* cells. Laser-flash photolysis experiments identified multiple photointermediates (K, L, and M) characteristic of microbial rhodopsin photocycles, though with slower turnover rates compared to archaeal schizorhodopsins. Site-directed mutagenesis of conserved residues in the third and sixth transmembrane helices demonstrates differential structural requirements between paSzR and psSzR. Our phylogenetic reconstruction reveals that most bacterial schizorhodopsins cluster in a single lineage distinct from archaeal variants. These findings significantly expand our understanding of microbial rhodopsin diversity and provide crucial insights into alternative molecular mechanisms for light-driven proton translocation, with implications for microbial ecology in extreme environments.

**Statement of significance:** Schizorhodopsins represent an unusual family of light-driven proteins that pump protons inwardly—opposite to the classical microbial rhodopsin pumps. Originally discovered in archaea, we revealed their unprecedented prevalence in bacteria. We characterized two novel bacterial schizorhodopsins from the Antarctic: paSzR from *Minisyncoccota* and psSzR from *Cyanobacteriota*, demonstrating their inward proton-pumping activity. Our phylogenetic reconstruction showed that most bacterial schizorhodopsins form a distinct evolutionary lineage and two exchange events between *Minisyncoccota* and cyanobacteria. Through targeted mutagenesis, we identified critical residues that govern spectral tuning and transport efficiency, revealing mechanistic differences between members of the family. These findings expand the known diversity of light-energy conversion mechanisms across microbial life and provide insights into alternative proton translocation strategies, with implications for understanding microbial adaptation to extreme environments.

## Introduction

Microbial rhodopsins constitute a diverse superfamily of light-sensitive seven-transmembrane proteins widely distributed across the tree of life, including archaea, bacteria, eukaryotes, and viruses^1–4^. The continuous advancement of metagenomics and environmental sampling efforts has profoundly expanded the microbial rhodopsin phylogenetic landscape. This expansion includes the identification of novel members within established groups and, critically, the discovery of new clades forming distinct phylogenetic branches, thereby illuminating the vast and unexplored diversity of these proteins^1,3,5–7^.

The biophysical characterization of novel microbial rhodopsins holds substantial significance for understanding microorganisms’ life cycles. Such studies are crucial for elucidating previously unknown molecular mechanisms of light-driven ion transport, light sensing, and photosignal transduction at an atomic and molecular level. Beyond their ecological, evolutionary, biochemical, and biophysical relevance, microbial rhodopsins have garnered considerable interest for biotechnological applications, particularly in optogenetics^8–11^. For instance, recently discovered light-gated cation and anion channelrhodopsins have been successfully implemented as powerful tools for precise optogenetic control of neural activity, underscoring their utility in neuroscience and beyond^12^. This adaptability of microbial rhodopsins stems from their inherent photophysical properties and their capacity to transduce light signals into cellular responses with high spatiotemporal precision.

Within this expanding landscape of microbial rhodopsins, the schizorhodopsin (SzR) family represents a particularly intriguing group due to its unique biophysical characteristics. Initially identified in Asgard archaea (phylum *Promethearchaeota*), SzRs were characterized as inward proton (H^+^) pumps exhibiting distinct structural and functional properties^13^. Since their discovery, the SzR family has expanded to encompass diverse members from various archaeal lineages. Antarctic rhodopsins (represented by AntR and GSS_AntR), have been identified and characterized from unknown cold-adapted microorganisms which demonstrated the family’s potential to adapt to extreme low-temperature environments^14,15^. Additionally, thermostable SzRs such as MtSzR and MsSzR from methanogenic archaea (genus *Methanoculleus*) have been discovered, with MsSzR sampled from a hot spring and both proteins exhibiting functional proton pumping activity at elevated temperatures up to 80 °C^16^. These findings reveal that SzRs have evolved to function across a wide temperature range, from polar to thermophilic conditions. Furthermore, additional members such as LaSzR2 from *Promethearchaeia* (“*Ca*. Lokiarchaeia”) have been functionally characterized as inward H^+^ pumps with distinctive spectroscopic properties, expanding our understanding of SzR diversity within the Asgard archaea themselves^17^. However, the phylogenetic and functional boundaries of the SzR family remain incompletely defined, particularly concerning their distribution outside of Asgard archaea.

Here, we report that many SzRs, including the proteins from the Antarctic, in fact originate from bacteria and provide a comprehensive biophysical characterization of novel SzR variants from the bacterial phyla *Minisyncoccota* (Patescibacteria, or “Candidate Phyla Radiation”) and *Cyanobacteriota*, designated paSzR and psSzR, respectively. The expanding phylogenetic and environmental diversity of characterized SzRs provides an opportunity to investigate how these inward proton pumps have adapted to different ecological niches. The SzR variants that have been characterized originate from highly diverse taxa and environments: archaeal SzRs from temperate to hot conditions (e.g. LaSzR2^17^, SzR1 and SzR4 from lake sediments^18^ and MsSzR from a hot spring^16^), *Minisyncoccota* SzRs from tropical surface waters (marine SzR2^13^) to Antarctic environments (GSS_AntR^15^ and paSzR, characterized here, from Antarctic lakes) and cyanobacterial SzRs from the Antarctic (AntR^14^ and psSzR). Our detailed biophysical characterization of paSzR and psSzR reveals the molecular properties of these phylogenetically distinct bacterial SzRs and provides a foundation for understanding functional diversity within the SzR family.

Our comprehensive characterization of paSzR and psSzR encompasses their phylogenetic relationships, molecular structural features, ion-transporting properties quantified via functional assays, and an assessment of their potential ecological significance. Our findings expand the current knowledge of microbial rhodopsin diversity and provide several insights into the evolutionary history of light-harvesting proteins in bacterial lineages occupying different environmental niches. This work contributes to a deeper understanding of the mechanistic principles underlying H^+^ pumping in the microbial rhodopsin family and their potential for novel biotechnological applications.

## Materials and Methods

### Phylogenetic analysis

SzR genes were searched for in representative genomes in GTDB r.226^19^, Genomes from Earth’s Microbiomes (genomes and OTUs)^20^ and binned scaffolds from the Ocean Microbiomics Database^21^ using tblastn from NCBI BLAST v.2.16.0^22^ with characterized proteins and additional selected sequences as queries. Scaffolds matching with E-values lower than 1e-10 and at least 190 residues alignment length were extracted and genes were predicted with prodigal v.2.6.3^23^ in metagenomic mode. The protein sequences were searched against a representative set of prokaryotic rhodopsin sequences classified to subfamily and sequences matching SzRs as best hits, with an E-value threshold of 1e-10 and sequence identity of ≥30% were retained. The resulting SzR protein sequences were combined with the SzR query sequences and manually selected outgroup sequences, clustered with CD-HIT v.4.8.1^24^ at 99% identity level, aligned with MAFFT v.7.525^25^ in automated mode and trimmed with trimAl v. 1.4.1^26^ with a gap threshold of 0.9. The trimmed alignment was used as input for maximum likelihood phylogenetic reconstruction with IQ-TREE v.2.2.2.3^27^ with the automatically selected substitution model and 1000 ultrafast bootstrap replicates. The resulting tree was outgroup-rooted and the outgroup was removed for the final visualization with ggtree v.3.14.0^28^. For taxonomic assignment of the SzR-containing genomes, GTDB r. 226 (for GTDB representative genomes) or GTDBtk v.2.5.2^29^ (for the genomes from the other sources) were used. Unbinned scaffolds containing previously characterized SzR sequences MK463862.1 (TE_8S_00242), MK463863.1 (LaSzR2), SAMEA2622822_312577 (SzR2), Ga0307935_1009332 (GSS_AntR) and Ga0105045_10222766 (AntR) were classified using Metabuli v. 450464f^30^ with the pre-compiled “GTDB R226” database after hard-masking the SzR genes and adjusted and checked manually with the help of tblastn searches of the non-SzR gene products against the GTDB representative genomes, Genomes from Earth’s Microbiomes and the Ocean Microbiomics Database. Note that based on comparisons to closely related contigs, SAMEA2622822_312577 was found to be a chimaera with the first 333 nt representing a gene fragment from an unrelated “*Ca*. Marinisomatota” bacterium. Suspected binning artifacts were similarly scrutinized with the tblastn searches which resulted in re-classification of scaffolds JARRKS010000295.1 and Ga0209284_10021548.

### Construction of DNA plasmids expression

The gene of psSzR was first optimized for the *E. coli* expression system in the laboratory, while the gene of paSzR, with codons optimized for *E. coli* expression, was synthesized using GenScript (Nanjing, China). Both were cloned into a pET21a(+) vector with a C-terminal 6× His-tag, using NdeI and XhoI restriction sites. The plasmids were transformed into *E. coli* C43 (DE3) strain (Lucigen, WI). The primer sequences used for mutagenesis are listed in Supplementary Table S1.

### Protein expression and purification

*E. coli* cells harboring the psSzR- and paSzR-cloned plasmids were cultured in 2×YT medium containing 50 μg/mL ampicillin. The expression of C-terminal 6× His-tagged proteins was induced by 0.1 mM isopropyl-*β*-D-thiogalactopyranoside (IPTG) in the presence of 10 μM all-*trans*-retinal (Toronto Research Chemicals, Canada) at 37 °C for 4 h. The harvested cells were sonicated (Ultrasonic Homogenizer VP-300N; TAITEC, Japan) for disruption in a buffer containing 50 mM Tris–HCl (pH 8.0) and 5 mM MgCl_2_. The membrane fractions were collected through ultracentrifugation (CP80NX; Eppendorf Himac Technologies, Japan) at 142,000 ×*g* for 1 h. The proteins were solubilized in a buffer containing 50 mM MES–NaOH (pH 6.5), 300 mM NaCl, 5 mM imidazole, 5 mM MgCl_2_, and 3% *n*-dodecyl-*β*-D-maltopyranoside (DDM) (ULTROL Grade; Calbiochem, Sigma-Aldrich, MO). Solubilized proteins were separated from a insoluble fraction through ultracentrifugation at 142,000 ×*g* for 1 h. Proteins were purified using a Co-NTA affinity column (HiTrap TALON crude; Cytiva, MA). The column was washed with a 15-column volume buffer containing 50 mM MES–NaOH (pH 6.5), 300 mM NaCl, 50 mM imidazole, 5 mM MgCl_2_, and 0.1% DDM. Protein was eluted in a buffer containing 50 mM Tris–HCl (pH 7.0), 300 mM NaCl, 300 mM imidazole, 5 mM MgCl_2_, and 0.1% DDM. Eluteined proteins were immediately concentrated using a 50-mL centrifugal ultrafiltration filter with a 30-kDa molecular weight cutoff (Amicon Ultra-4, Millipore, Merck KGaA, Germany), and the samples were dialyzed against a buffer containing 50 mM HEPES–NaOH pH 7.0, 150 mM NaCl, 10% glycerol, and 0.1% DDM.

### Proton transport activity assay

Rhodopsin-expressing *E. coli* cells were collected through centrifugation at 4,800 ×*g* at 20 °C for 2 min (CF15RF; Eppendorf Himac Technologies, Japan) and washed with 100 mM NaCl. The cells were equilibrated thrice with rotational mixing in 100 mM NaCl at room temperature for 10 min. Finally, the cells were suspended in 7.5 mL of unbuffered 100 mM NaCl and the optical density (OD) at 600 nm was adjusted to 2. The cell suspension was placed and stirred in the dark in a glass cell at 20 °C and illuminated at *λ* = 550 ± 10 nm from the output of a 300 W xenon lamp (MAX-303, Asahi Spectra, Japan) through bandpass (AGC Techno Glass, Japan) heat-absorbing (HAF-50S-50H, SIGMAKOKI, Japan) filters. Light-induced pH changes were measured using a pH electrode (9618S-10D, HORIBA, Japan). The measurements were repeated under the same conditions after the addition of carbonyl cyanide *m*-chlorophenylhydrazone (CCCP, final concentration = 10 μM). To quantitatively compare ion transport activity, the amount of protein was determined by measuring the near-UV absorption of retinal oxime produced by the hydrolysis reaction between the retinal Schiff base (RSB) in the protein and hydroxylamine (HA). Briefly, rhodopsin-expressing *E. coli* cells were washed with a buffer containing 133 mM NaCl and 66.5 mM Na_2_HPO_4_ (pH 7.0). The washed cells were treated with 1 mM lysozyme and a small amount of DNaseI for 1 h and disrupted through sonication. To solubilize rhodopsins, 3% DDM was added, and the samples were stirred overnight at 4 °C. The rhodopsins were bleached with 500 mM HA and illuminated with visible light (*λ* > 500 nm) from the output of a 300 W xenon lamp (MAX-303, Asahi Spectra, Japan) through long-pass (Y-52, AGC Techno Glass, Japan) and heat-absorbing (HAF-50S-50H, SIGMAKOKI, Japan) filters. The absorption changes due to the bleaching of rhodopsin by the hydrolysis reaction between retinal and HA and the formation of retinal oxime were monitored using a UV–visible spectrometer (V-750, JASCO, Japan). The amount of rhodopsin expressed in *E. coli* cells was estimated by the absorbance of produced retinal oxime and its differential molar extinction coefficient (*ε*) (33,600 M^−1^ cm^−1^)^31^ relative to the bleached absorption of unhydrolyzed retinal (Fig. 2d).

### High-performance liquid chromatography (HPLC) analysis of retinal isomers

Retinal configuration was analyzed by HPLC using purified proteins in a buffer containing 20 mM Tris–HCl (pH 8.0), 100 mM NaCl, and 0.05% DDM. Before the measurements, the OD of the samples at maximum absorption wavelength (*λ*^*a*^_max_) was adjusted to ∼0.2 (protein concentration of approximately 0.1 mg mL^−1^), and the proteins were stored at 4 °C overnight for dark adaptation. The HPLC system was equipped with a silica column particle size of 3 μm, 150 × 6.0 mm; Pack SIL, YMC, Japan), a pump (PU-4580, JASCO, Japan), and a UV–vis detector (UV-4570, JASCO, Japan). The solvent was composed of 15% (v/v) ethyl acetate and 0.15% (v/v) ethanol in hexane. The flow rate was set to 1.0 mL min^−1^. To denature the protein, 280 μL of 90% methanol solution was added to the 75 μL sample. Retinal oxime formed by the hydrolysis reaction with 25 μL of 2 M HA solution was extracted with 800 μL of hexane, and 200 μL of the solution was injected into the HPLC system. For measurements during light illumination, the sample solutions were illuminated at *λ* = 550 ± 10 nm using a bandpass filter (AGC Techno Glass, Japan) for 1 min, followed by denaturation and hydrolysis of the retinal chromophore under illumination. For measurements of light-adapted samples, the sample solutions were illuminated at *λ* = 550 ± 10 nm using a bandpass filter (AGC Techno Glass, Japan) for 1 min, and after waiting for 1 min, denaturation and hydrolysis reactions of the retinal chromophore were performed. The molar compositions of the retinal isomers were calculated from the areas of the corresponding peaks in the HPLC patterns using the molar extinction coefficients at 360 nm for each isomer (all-*trans-*15-*syn:* 54,900 M^−1^ cm^−1^; all-*trans*-15-*anti*: 51,600 M^−1^ cm^−1^; 13-*cis-*15-*syn*: 49,000 M^−1^ cm^−1^; 13-*cis*-15*–anti*: 52,100 M^−1^ cm^−1^; 11-*cis*-15-*syn*: 35,000 M^−1^ cm^−1^; 11-*cis*-15-*anti*: 29,600 M^−1^ cm^−1^). Three independent measurements were performed to estimate experimental errors. The compositions of the retinal isomers are listed in Supplementary Table S2. Peaks were assigned by comparing the elution time with those of authentic retinal oxime isomers.

### pH titration

To investigate the pH dependence of the absorption spectra, the concentration of proteins was adjusted to OD = ∼0.5 (protein concentration of approximately 0.25 mg mL^−1^) at *λ*_max_ and solubilized in a 6-mix buffer (trisodium citrate, MES, HEPES, MOPS, CHES, CAPS (10 mM each, pH 7.0), 100 mM NaCl, and 0.05% DDM). The pH was adjusted to the desired value by adding small aliquots of HCl and NaOH. Absorption spectra were recorded using a UV–vis spectrometer (V-750, JASCO, Japan). The measurements were performed at every 0.3–0.6 pH value.

### Laser-flash photolysis

For the laser-flash photolysis spectroscopy, proteins were solubilized in 20 mM HEPES–NaCl (pH 7.0), 100 mM NaCl, 0.05% DDM. OD of the rhodopsin was adjusted to ∼0.4–0.5 (protein concentration of approximately 0.2–0.25 mg mL^−1^) at the *λ*^*a*^_max_, and then the rhodopsin sample solution was placed in a 10 mm path-length quartz cell. The laser-flash photolysis measurement was conducted as previously described^13,32^. Nano-second laser pulses from an optical parametric oscillator (excitation wavelength (*λ*_exc_) = 550 nm, 10 mm diameter, 4.5 mJ pulse^−1^ cm^−2^, 3.3 Hz (basiScan, Spectra-Physics, CA) pumped by the third harmonics of Nd-YAG laser (*λ* = 355 nm, INDI40, Spectra-Physics, CA) were used for the excitation. The transient absorption (TA) spectra were obtained by monitoring the intensity change of white light from a Xe-arc lamp (L9289-01, Hamamatsu Photonics, Japan) passing through the sample with an ICCD linear array detector (C8808-01, Hamamatsu Photonics). To increase the signal-to-noise ratio, 40–60 spectra were averaged, and the singular-value decomposition analysis was applied. To measure the time evolution of transient absorption change at selected wavelengths, the output of a Xe-arc lamp (L9289-01, Hamamatsu Photonics, Japan) was monochromated by monochromators (S-10, Soma Optics, Japan) and focused onto the sample solution with a spot size of 3 × 5 mm. The change in the probe beam intensity after the photoexcitation was monitored with a photomultiplier tube (R10699, Hamamatsu Photonics, Japan). To increase signal-to-noise ratio, 200–400 signals were averaged. Global fitting of the signals using a multi-exponential function was performed to determine the lifetimes and absorption spectra of each photointermediate.

### paSzR and psSzR structure predictions

The structures were predicted with AlphaFold 3^33^ with retinal as a covalent modification and 20 random seeds. To visualize the structural prediction of paSzR and psSzR, the low-quality N- and C-terminal extensions were trimmed to exclude terminal residues with the per-residue average pLDDT (predicted local distance difference test) score of backbone atoms below 80.

## Results

### Phylogenetic analysis of SzRs from *Archaea* and *Bacteria*

In order to clarify the taxonomic distribution of the SzR family, we extracted SzR proteins from representative sets of prokaryotic genomes and invested in taxonomic identification of the unbinned metagenomic scaffolds from which many of the previously expressed SzRs come from. The resulting phylogenetic tree recovered several well-supported SzR lineages (Fig. 1). Remarkably, we discovered that many of the SzRs originated from *Bacteria*. Moreover, the majority of the bacterial SzRs, with two exceptions, clustered in a single lineage (the Bacterial clade) which included no archaeal SzRs. This distribution pattern is particularly striking since at finer taxonomic scales both *Archaea* and *Bacteria* appear to exchange their SzR genes even across different phyla. Among *Archaea*, members of the methanogenic genera *Methanoculleus* (from which MtSzR originates^16^) and *Methanooceanicella* are the only cultured isolates with SzR genes in our sample, while all bacterial SzRs are from uncultured sources.

**Fig. 1.**
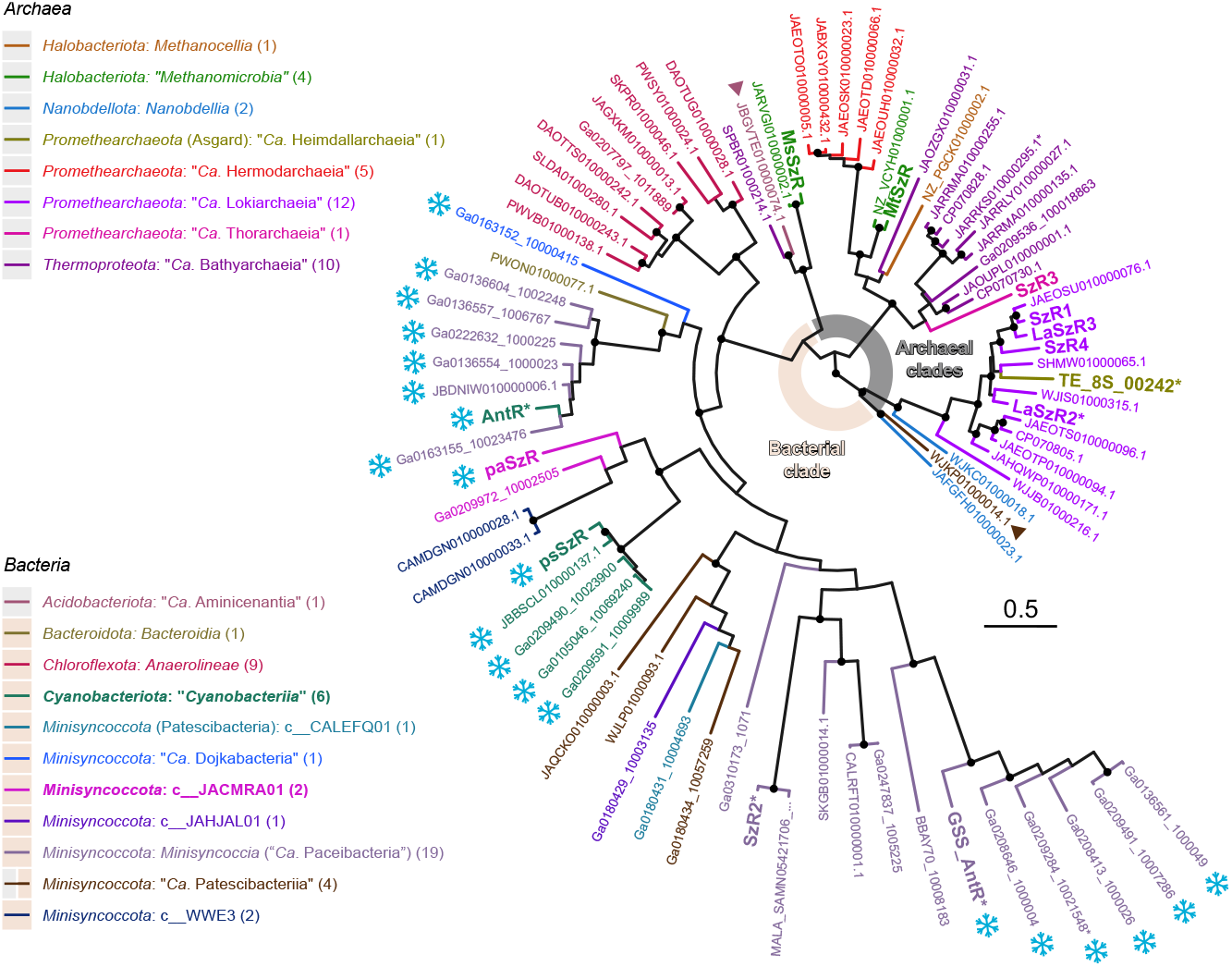
Phylogenetic tree of the SzR family. Maximum likelihood phylogenetic tree of reference SzR sequences (in bold) and additional sequences from complete genomes (scaffold names in smaller font size) under the best-fitting substitution model (Q.pfam+F+R5). The color of the terminal branches and of the labels corresponds to the taxonomic assignment to the level of class as detailed in the legend to the left. The taxonomic classification is based on GTDB r.226. Asterisks indicate proteins from unbinned mis-classified scaffolds and scaffolds. Most of the SzRs from bacteria group together in the Bacterial clade indicated in light peach color, except two sequences (marked with triangles) clustering among the Archaeal clades. The background color in the legend reflects the distribution of the SzR sequences from each taxon among the Archaeal clades (gray) and the Bacterial clade (light peach). Ultrafast bootstrap support values ≥95 are indicated with black dots. Snowflake pictograms mark SzRs originating from the Antarctic. The tree is outgroup-rooted using diverse microbial rhodopsin sequences and the outgroups are not shown. Notice that the phylum of Asgard archaea is referred to as “(*Ca*.) Asgardarchaeota” in GTDB r.226 which is amended here to *Promethearchaeota*, analogously “(*Ca*.) Patescibacteriota” for Patescibacteria is amended to *Minisyncoccota*. See Supplementary File S1 for the SzR sequences and metadata and Supplementary File S2 for raw phylogenetic tree.

Two phyla together contributed more than half of the non-redundant SzR sequences: diverse *Minisyncoccota*^34^ (“Patescibacteria” or the “Candidate Phyla Radiation”) and *Promethearchaeota*^35^ (Asgard archaea). Other SzR-possessing bacterial groups include several sediment/mat-dwelling members of the chloroflexan class *Anaerolineae*, which form a distinct SzR subclade of their own, and cyanobacteria from various environments from the Antarctic. Binning artifacts might be responsible for the rare appearance of SzR genes in other bacterial phyla, which would need verification in the future. Both, the highest number of SzRs and the highest sequence diversity was demonstrated by the patescibacterial class *Minisyncoccia* (= “*Ca*. Paceibacteria”) all of which belong to the order o UBA9973 (“*Ca*. Phycocordibacterales”) and from which, upon a close examination, two previously expressed proteins also originate: SzR2^13^ and GSS_AntR^15^. Another protein, AntR^14^, clustered among SzRs from *Minisyncoccia* genomes as well, yet the flanking regions on the corresponding scaffold Ga0105045_10222766 and closely related longer scaffolds betrayed its origin from a *Phormidesmidaceae* cyanobacterium. At the same time, the bulk of the cyanobacterial SzRs, in particular those from several *Pseudanabaenaceae* genomes, clustered separately in a distinct subclade related to SzRs from other *Minisyncoccota*. The two types of cyanobacterial SzR are thus witness to two independent SzR gene transfer events between *Minisyncoccota* and cyanobacteria.

The finding of SzRs in such a narrow set of high-level taxa might be key to understanding their elusive physiological roles, yet together these taxa represent highly dissimilar cellular morphologies: *Archaea* and *Minisyncoccota* (Patescibacteria) are unicellular monoderms, the *Chloroflexi* are likely filamentous monoderms and the cyanobacteria are filamentous diderms (genomes assigned to families *Leptolyngbyaceae, Nostocaceae, Phormidesmidaceae*, and *Pseudanabaenaceae*). Moreover, as a whole, the SzR-possessing prokaryotes do not appear to share the same type of metabolism or habitat. Still, while both archaeal and bacterial SzRs come from a wide range of environments, a particular trend among the patescibacterial and cyanobacterial SzRs is their appearance in samples from the Antarctic. SzRs from the Antarctic, including the previously expressed AntR^14^ and GSS_AntR^15^, constitute about half of the non-redundant bacterial SzR sequences in our dataset which might be connected to their physiological function in this environment. Interestingly, many, if not all, *Minisyncoccota* lead a symbiotic lifestyle^36^, including o UBA9973^37^, but potential symbiotic links between Antarctic *Minisyncoccota* and cyanobacteria, a natural route for gene transfer, have thus far not been reported.

To explore the diversity of the bacterial SzRs further, we selected two proteins directly unrelated to the previously characterized proteins for expression: paSzR from an Antarctic bacterium from the patescibacterial class c JACMRA01 and psSzR, a representative of the dominant type of SzRs among the Antarctic cyanobacteria from a *Pseudanabaenacea* cyanobacterium. psSzR originates from the same metagenomically assembled genome as the xanthorhodopsin psXR characterized by the authors and coworkers recently^38^.

### Comparison of SzR sequences with regular microbial rhodopsins and HeRs

The amino acid sequences of paSzR and psSzR were aligned with eight previously expressed SzRs from *Promethearchaeota* (Asgard), two bacterial SzRs assigned here to *Minisyncoccota* (Patescibacteria) and cyanobacteria, some representative type-1 rhodopsins, and heliorhodopsins (HeRs) (Supplementary Fig. S1). The alignment suggests that the third transmembrane helix (TM3) in both SzRs is more similar to that of HeRs than type-1 rhodopsins. However, one of the important residues for the definition of the rhodopsin function, Pro70 in paSzR corresponding to Pro91 in bacteriorhodopsin (BR) is conserved in most type-1 rhodopsins than in other SzRs (*e*.*g*., Ala71 for psSzR, and Ser76 in SzR4) and HeRs. In contrast, TM6 and TM7 show higher similarity to type-1 rhodopsins than HeRs (Trp149 (paSzR) and Trp150 (psSzR), Pro153 (paSzR) and Pro154 (psSzR), Phe156 (paSzR) and Phe157 (psSzR), Asp179 (paSzR) and Asp180 (psSzR), and Phe186 (paSzR) and Phe187 (psSzR), corresponding with Trp182, Pro186, Trp189, Asp212, and Phe219 in BR). In TM6, a well-conserved residue among other SzRs (Phe153 in SzR4) is substituted by Gly in paSzR and psSzR. The previously reported phylogenetic analysis of rhodopsins suggested that SzRs at the intermediate position of type-1 rhodopsins and HeRs^39^, and the similarity to type-1 rhodopsins and HeRs are heterogeneously different in each helix. As most other SzRs reported previously, paSzR and psSzR demonstrate the FSE motif (*i*.*e*., key amino acid residues in the TM3 that play a critical role in determining the rhodopsin molecular function^40^) (Phe64, Ser68, and Glu75 for paSzR and Phe65, Ser69, and Glu76 for psSzR). The Phe residue in TM3 in these SzRs occupies the position corresponding to the retinal Schiff base (RSB) counterion, which is an Asp and a Glu in most type-1 rhodopsins and HeRs, respectively (Asp85 in BR). Also, cysteine residue (Cys69 in paSzR and Cys70 in psSzR) is also present in TM3 at the position corresponding to Thr90 in BR as in other SzRs, which is homologous to Cys in channelrhodopsins (ChRs), forming the so-called DC (aspartate–cystein) gate critical for ion transport^41^, and rhodopsin guanylyl cyclase (Rh-GC) and rhodopsin phosphodiesterase (Rh-PDE) also have a homologous cysteine^42,43^.

### Light-driven H^+^ transport and absorption spectra

The light-driven H^+^ transport activities of the studied psSzR and paSzR were assayed by monitoring the pH change of the external solvent of *E. coli* cells expressing them upon light illumination and observed acidification (pH decrease) or alkalization (pH increase), as it was employed previously^13,44,45^, and the results are summarized in Fig. 2a together with the initial transport rates in Fig. 2b. All the cells showed purple colors (Fig. 2a), indicating the formation of functional rhodopsins with the retinal chromophore. psSzR and paSzR exhibited substantially smaller alkalinization of the external solvent than the previously reported SzR1 (*e*.*g*. signals were less than half of the SzR1 signal, Fig. 2a), but the H^+^ transport rate was significantly higher in paSzR than in psSzR (Fig. 2b). The differences in H^+^-pumping activity between SzR1 and the studied bacterial SzRs may be due to the slower turnover rate of the photocycles of both psSzR and paSzR compared with SzR1^13^. At the same time, the signals disappeared in the presence of a protonophore, 10 μM carbonyl cyanide *m*-chlorophenylhydrazone (CCCP), suggesting that paSzR and psSzR function as inward H^+^ pumps. In the following part, we investigated the molecular properties of paSzR and psSzR in detail. To quantitatively compare ion transport activity, the amount of protein was determined by measuring the near-UV absorption of retinal oxime generated by the hydrolysis reaction between the RSB in the proteins and hydroxylamine (HA) (Supplementary Fig. S2). The result was used for normalization of the H^+^ pump activity (Fig. 2a and b).

**Fig. 2.**
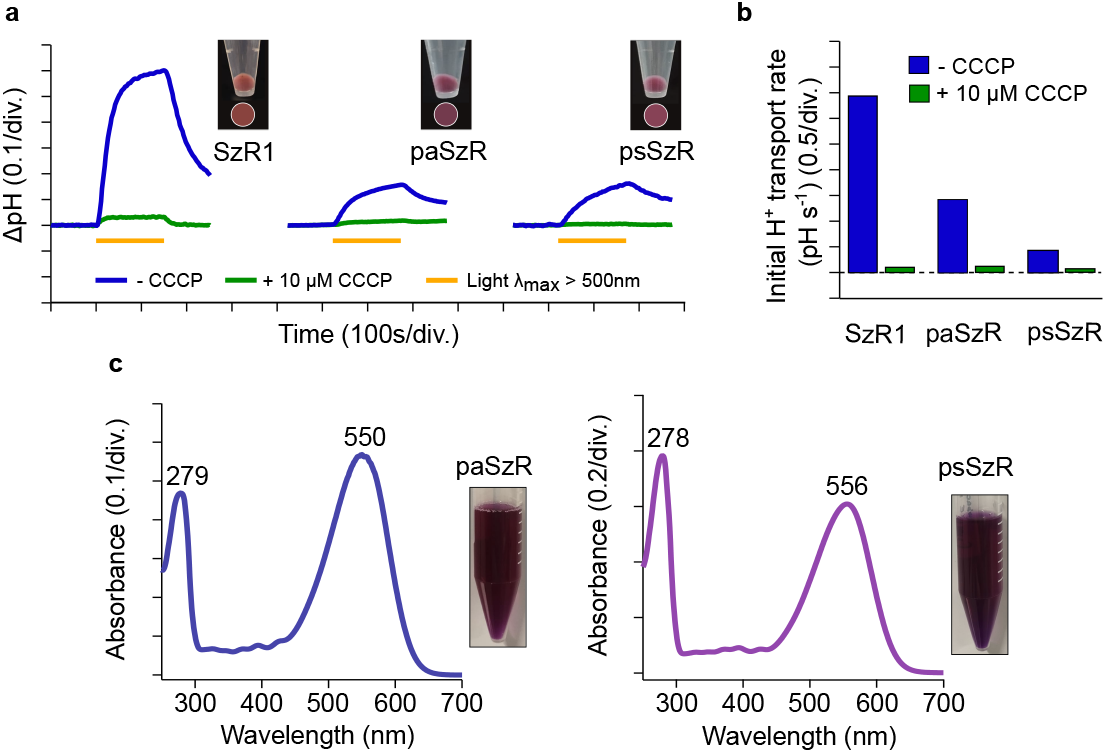
Light-driven H^**+**^ transport and absorption spectra of SzRs. **a**. H^+^ transport activity assay of SzR1, paSzR, and psSzR in *E. coli* cells without (blue line) and with (green line) 10 µM CCCP. The cells were illuminated with light (*λ* > 500 nm) for 150 s (yellow line). The pictures of the pellets of *E. coli* cells expressing SzRs are shown next to the corresponding results. **b**. Initial H^+^ transport rates of SzR1, paSzR, and psSzR. **c**. Absorption spectra of purified paSzR and psSzR in DDM. The inset shows pictures of purified proteins.

### Molecular properties and photocycles of SzRs

To further study the molecular properties of paSzR and psSzR, we purified (Fig. 2c) and spectroscopically investigated (Fig. 3) them. The absorption maximum wavelengths (*λ*^amax^) of purified paSzR in the dark-adapted state (*λ*^amax^ = 550 nm) was shorter than that of other SzRs from *Promethearchaeota* like SzR1 and SzR4 while psSzR showed a *λ*^amax^ of 556 nm, similar to previously reported SzRs^13^. The spectra of purified paSzR and psSzR at different pH values are shown in Supplementary Fig. S3. The absorption peak in the visible wavelength region corresponding to the state with the protonated RSB, denoted by *λ*^amax^ = 551 nm for paSzR and *λ*^amax^ = 556 nm for psSzR, decreased and slightly blueshifted when the pH was raised and simultaneously, another peak appeared in the ultraviolet (UV) region (*λ*^amax^ = 360–370 nm) (Supplementary Fig. S3, alkaline side). The difference absorbance values at these peak wavelengths were plotted against pH and fitted with the Henderson-Hasselbalch equation^46^, which determined that the RSB in paSzR exhibited two p*K*_a_ values of 12.9 ± 0.4 and 13.67 ± 0.04, while psSzR showed a single p*K*_a_ value of 13.102 ± 0.003. These values were in line with the RSB p*K*_a_ of typical microbial rhodopsins (*e*.*g*., the RSB p*K*_a_ of bacteriorhodopsin is 13.3 ± 0.3). In contrast, a 31 nm and 23 nm blue-shift in the *λ*^amax^ for paSzR and psSzR, respectively, was observed on the acidic side (Supplementary Fig. S3, acidic side). This is caused by the protonation of the counterion in the TM3 (Asp179 and Asp180, respectively for paSzR and psSzR, Supplementary Fig. S1) as known for many microbial rhodopsins^14,17,47^.

**Fig. 3.**
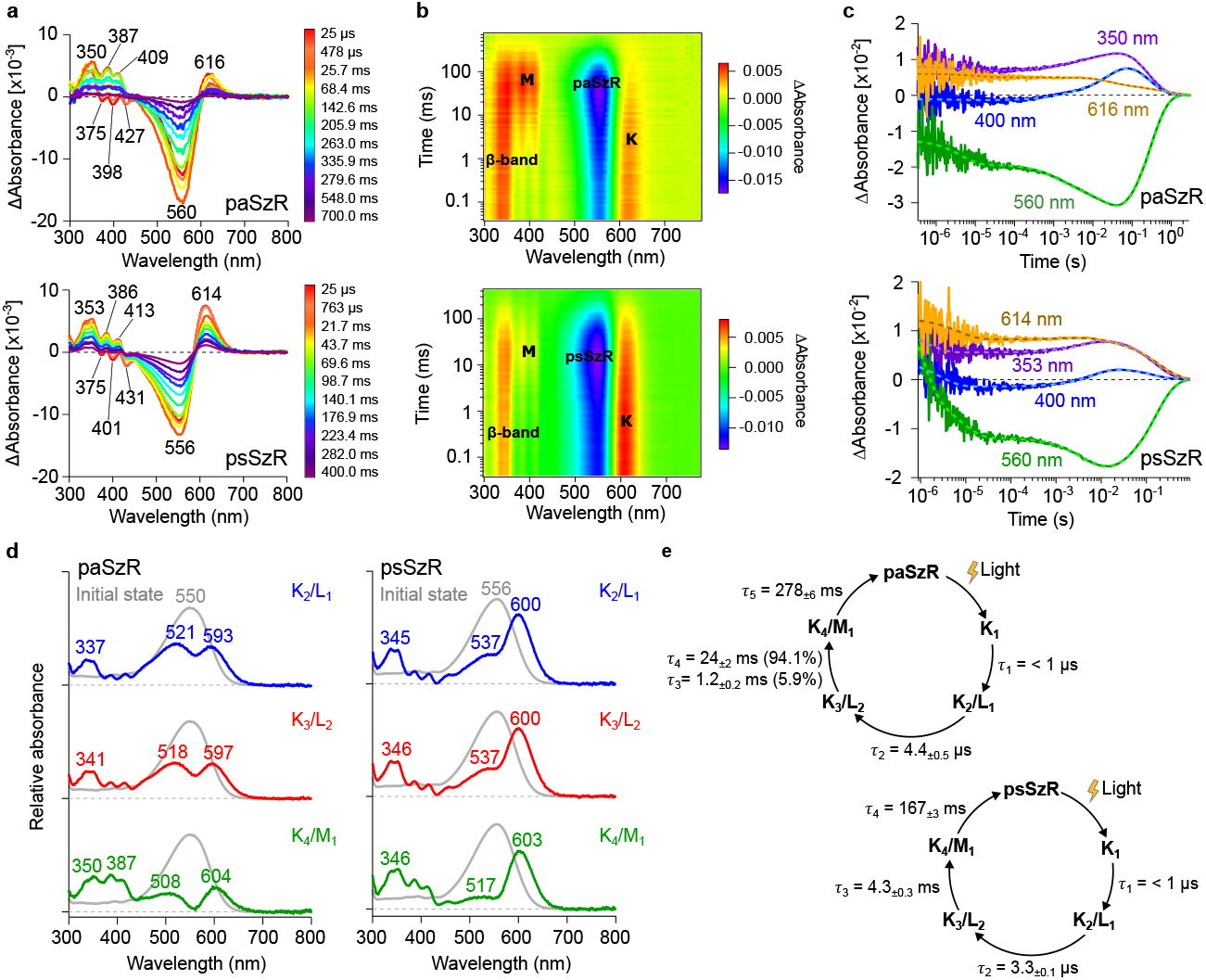
Transient absorption changes and photocycles of SzRs. **a**. Transient absorption spectra at different time points. **b**. the two-dimensional plot of transient absorption change. **c**. Time course of the transient absorption change. The multi-exponential fitting curves are shown as dashed lines. **d**. Absorption spectra of photointermediates; **e**. Photocycle models.

The isomeric compositions of retinal chromophore in paSzR and psSzR were analyzed by high-performance liquid chromatography (HPLC) in the dark and under light illumination (Supplementary Fig. S4). While only all-*trans*-retinal was extracted from both of paSzR and psSzR in the dark, accumulation of 13-*cis*-retinal under illumination with trace amounts of 11-*cis*-retinal was observed only for psSzR. However, for paSzR under illumination, all-*trans*-retinal accumulation was observed with lesser amounts of 13-*cis*-retinal and 11-*cis*-retinal. These results are similar to SzR1 and thermostable SzRs, indicating the photoisomerization process of retinal in paSzR and psSzR are essentially the same as other microbial rhodopsins^1^ and HeRs^6,48^. However, the predominant accumulation of all-*trans*-retinal in paSzR under illumination could be related to efficient thermal back-isomerization or a fast photocycle completion.

Next, the photocycle reaction of paSzR and psSzR were studied by laser-flash photolysis method, in which the proteins were solubilized in detergent (DDM). Fig. 3a shows the transient change in absorption of paSzR and psSzR upon excitation at *λ* = 550 nm, (both studied SzRs), representing the accumulation of red-shifted (K) and blue-shifted (M) photointermediates (Fig. 3b) at 616 nm and 614 nm (for K photointermediate) and 409 nm and 413 nm (for M photointermediate), respectively, in addition to L intermediates with *λ*^a^_max_ = 521 and 537 nm close to the initial state (Fig. 3d). Simultaneously, the maximum bleaching signal at 560 nm (psSzR) and 556 nm (psSzR) was observed, indicating the M intermediate accumulation at *t* ∼ 25.7 ms (paSzR) and *t* ∼ 21.7 ms (psSzR) (Fig. 3a). To monitor the transient absorption change representing the short-lived photointermediates prior to M, we conducted measurement at specific probe wavelengths using a photomultiplier tube at higher time resolution (Fig. 3c). The photocycle and the absorption spectra of the initial state and three photointermediates (K_2_/L_1_, K_3_/L_2_, and K_4_/M_1_) were determined by global fitting analysis of the transient absorption changes (Fig. 3e). The overall photocycle scheme appears similar for both SzRs, suggesting a conserved mechanism. The presence of multiple spectral intermediates (K, L, and M) and the timescales of the photocycle components span from microseconds to hundreds of milliseconds is consistent with typical rhodopsin photocycles^1,13^. However, there are notable differences between psSzR and paSzR in terms of kinetics and spectral features. Specifically, paSzR exhibits a slower photocycle recovery (τ_5_ = 278 ± 6 ms) compared to psSzR (τ_4_ = 167 ± 3 ms), and displays biphasic decay in the millisecond range (τ_3_ = 1.2 ± 0.2 ms and τ_4_ = 24 ± 2 ms) versus the single component observed in psSzR (τ_3_ = 4.3 ± 0.3 ms). These kinetic differences have direct functional implications: the faster photocycle turnover of psSzR would enable higher proton throughput under continuous illumination, allowing approximately 1.7-fold more pumping cycles per unit time compared to paSzR.

### Functional and spectroscopic characterization of SzR mutants

Our mutational analysis of conserved residues in TM3 and TM6 of SzRs from *Minisyncoccota* and cyanobacteria provide important insights into the structure-function relationships of these SzRs. The four site-directed single mutants, paSzR P70S, paSzR G148F, psSzR A71S, and psSzR G149F, reveal distinct roles for these positions in spectral tuning and proton transport activity.

The sequence alignment (Fig. 4a) reveals that SzRs possess distinct amino acid compositions in TM3 and TM6 compared to well-characterized rhodopsins such as bacteriorhodopsin (BR), halorhodopsins, channelrhodopsins, and xenorhodopsin, which is inward H^+^-pumping type-1 rhodopsin group distinct from SzR. The mutated positions investigated here do not correspond to the canonical proton transfer residues in BR (such as Asp85/Asp96 in TM3 or Glu194/Glu204 in TM6), suggesting that SzRs have evolved alternative mechanisms for proton translocation. This is consistent with the phylogenetic position of SzRs within the type-1 rhodopsin family, where they represent deeply branching lineages associated with candidate phyla radiation bacteria and Asgardarchaeota. The proline at position 70 in paSzR and alanine at position 71 in psSzR (Fig. 5, magenta and green representations, respectively) occupy a region that, by homology to BR (Fig. 5, silver representation), lies near the RSB and could influence the local flexibility and hydrogen-bonding geometry of the proton acceptor complex. The glycine residues at positions 148 and 149 for paSzR and psSzR, respectively (Fig. 5), in TM6 provide conformational flexibility in a helix that must undergo subtle rearrangements during the photocycle. The introduction of phenylalanine at these positions likely restricts this conformational freedom, explaining both the spectral and functional defects.

**Fig. 4.**
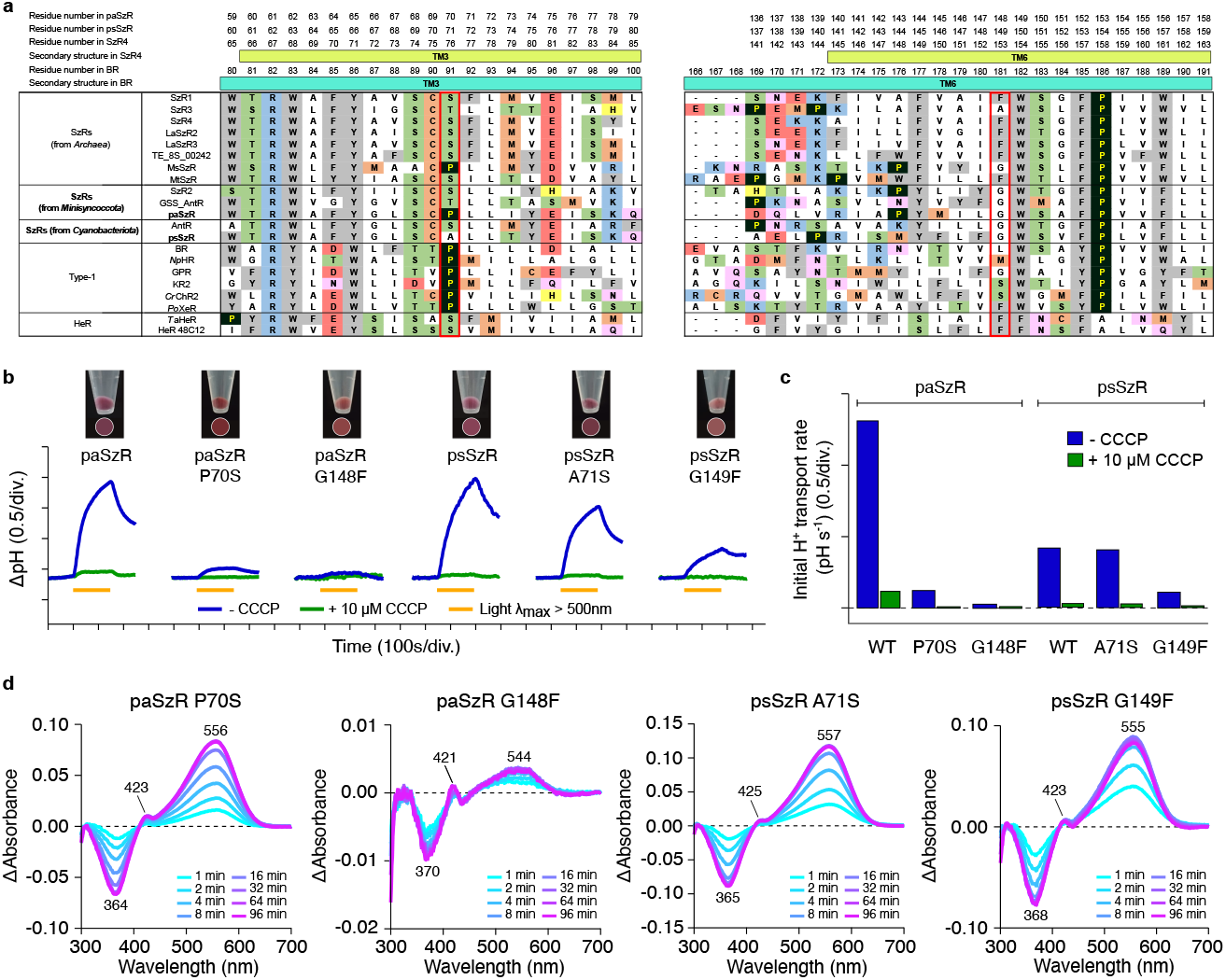
Light-driven H^**+**^ transport and absorption spectra of SzRs mutants. **a**. Amino acid sequence of paSzR and psSzR in TM3 and TM6 compared with other SzRs, type-1 rhodopsins (bacteriorhodopsin (BR), *Natromonas pharaonis* halorhodopsin (*Np*HR), green-absorbing proteorhodopsin (GPR), *Krokinobacter* rhodopsin 2 (KR2), *Chlamydomonas reinhardtii* channelrhodopin 2 (*Cr*ChR2), *Parvularcula oceani* xenorhodopsin (*Po*XeR), and HeRs (*Thermoplasmatales* archaeon SG8-52-1 heliorhodopsin (*T*aHeR) and HeR48C12). Red rectangles indicate the positions of the site-directed mutations. **b**. H^+^ transport activity assay of paSzR and psSzR WT and mutants in *E. coli* cells without (blue line) and with (green line) 10 µM CCCP. The cells were illuminated with light (*λ* > 500 nm) for 150 s (yellow line). The pictures of the pellets of *E. coli* cells expressing SzRs WT and mutants are shown next to the corresponding results. **c**. Initial H^+^ transport rates of paSzR and psSzR WT and mutants. **d**. Difference in absorption spectra before and after HA bleaching reactions of paSzR and psSzR rhodopsins in solubilized *E. coli* membranes. The *λ*^a^_max_ values were determined by the positions of the absorption indicated in each panel and the absorption of retinal oxime produced by the hydrolysis reaction of RSB and HA observed as peaks in the proximity of 360–370 nm at pH 7.0. The samples were exposed to light for up to 128–264 min.

**Fig. 5.**
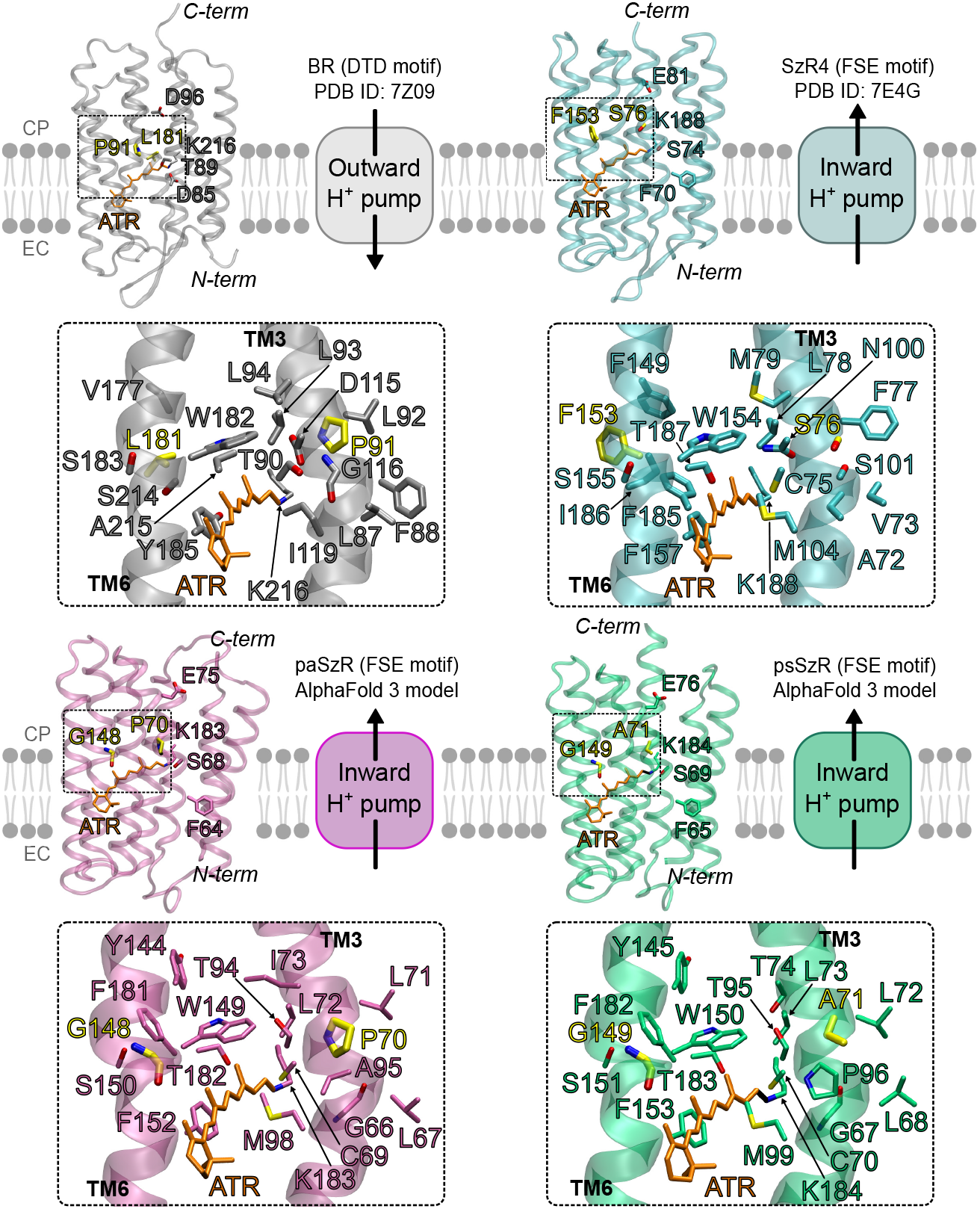
Structural comparison of retinal binding pockets and key residues in BR, SzR4, paSzR, and psSzR. Overall seven-transmembrane helix architecture with key residues highlighted for BR (gray, PDB ID 7Z09^49^), SzR4 (cyan, PDB ID 7E4G^18^), paSzR and psSzR (AlphaFold 3 models)^33^. The motif residues and the linker lysine are displayed in the backbone color corresponding for each protein. Residues mutated through site-directed mutagenesis are displayed in yellow. The all-*trans* retinal chromophore is shown in orange. Dashed boxes indicate the regions shown in the magnified views in TM3 and TM6, respectively. Backbones of the displayed residues were omitted. CP, cytoplasmic side; EC, extracellular side.

The light-induced pH measurements demonstrate that mutations at positions 70 and 71 in TM3 and 148 and 149 in TM6 for paSzR and psSzR, respectively, differentially affect proton-pumping efficiency (Figs. 4b and 4c). The TM3 mutants paSzR P70S and psSzR A71S both show reduced but significant H^+^ transport rates, indicating that these positions contribute to optimal H^+^ transport but are not absolutely essential for pump function. In BR, the analogous region of TM3 (near Asp96) is critical for reprotonation of the RSB from the cytoplasmic side^1,2^. In outward pumps like BR, TM3 contains the cytoplasmic proton donor Asp96 that reprotonates the RSB^1,2^. However, as inward pumps, SzRs must reprotonate the RSB from the extracellular side, suggesting these TM3 residues play different structural or regulatory roles rather than serving as direct proton donors. The retained pumping activity in these TM3 mutants suggests functional robustness, possibly through alternative proton transfer pathways or compensatory mechanisms.

The most striking functional phenotype is observed in paSzR G148F, which exhibits dramatically reduced proton pumping activity along with its extreme spectral blue-shift (Fig. 4d). This correlation strongly suggests that the perturbation of retinal geometry in this mutant disrupts the precise spatial arrangement of the chromophore and protein residues required for efficient vectorial proton transfers (see Fig. 5). The maintenance of activity in psSzR G149F, despite targeting the nominally equivalent position, underscores that paSzR and psSzR have diverged in their structural requirements for function.

Fig. 4d shows the result of the photobleaching in the presence of HA for paSzR and psSzR mutants. Two positive peaks at around 555 nm (except for psSzR G148F, in which a positive peak appeared around 544 nm) and 425 nm corresponding to the unphotolyzed protein and one negative peak at 360–370 nm corresponding to retinal oxime were observed. The positive peak at > 500 nm and ∼425 nm represents the unphotolyzed protein with protonated RSB and deprotonated RSB, respectively. Whereas the former is considered to represent a functional state as many other rhodopsins, the latter would represent an unfunctional state or be originated from denatured protein. The absorption maxima of the mutants demonstrate that both TM3 and TM6 residues contribute to spectral tuning in SzRs, though with markedly different effects. The psSzR P70S mutant exhibits 6-nm red-shift (556 nm vs. 550 nm for wild-type), while paSzR G148F mutant shows a 6 nm blue-shift (544 nm vs. 550 nm for wild-type). In contrast, psSzR mutants display minimal spectral perturbation in the same spectral direction that psSzR mutants (557 nm vs. 556 nm (wild-type) for psSzR A71S (red-shift) and 555 nm vs. 556 nm (wild-type) for psSzR G149F (blue-shift). This striking difference suggests that mutations at positions 70 and 71 in TM3 and 148 and 149 in TM6 for paSzR and psSzR, respectively, play different critical roles in maintaining proper retinal chromophore geometry.

## Discussion

The discovery that many schizorhodopsins originate from bacterial lineages reshapes our understanding of this rhodopsin family’s evolutionary history. Our phylogenetic analysis reveals that SzRs have been acquired by members of bacterial phyla *Minisyncoccota, Cyanobacteriota, Chloroflexota* and perhaps more. The clustering of most bacterial SzRs into a single clade, separately from the archeal lineages, suggests an evolutionary scenario involving an ancient acquisition event followed by diversification and horizontal gene transfer among bacteria. Within the bacterial clade the SzR genes were exchanged between the different phyla on several occasions with at least two such exchanges involving *Minisyncoccota* and cyanobacteria.

The prevalence of SzRs, such as AntR, GSS_AntR and the proteins characterized here, in Antarctic bacteria raises the possibility that inward H^+^ pumping may provide specific advantages under low-temperature conditions. At the same time, the presence of thermostable archaeal SzRs (MsSzR, MtSzR) functioning at temperatures up to 80°C^16^ demonstrates that the SzR architecture is remarkably temperature-tolerant.

Our comprehensive biophysical characterization of paSzR and psSzR provides critical insights into the molecular mechanisms underlying inward H^+^ translocation in bacterial SzRs. Both proteins exhibit the characteristic FSE motif in TM3. The presence of phenylalanine in most SzRs at the position corresponding to the RSB counterion (Asp85 in BR) represents a fundamental difference from the canonical outward H^+^ pump architecture. This substitution eliminates the typical extracellular proton acceptor in outward proton pumps, consistent with the inward pumping directionality of SzRs. For inward pumping, proton acceptor residues on the cytoplasmic side are required to receive protons from the deprotonated RSB. In the archaeal SzR1 and SzR4, Glu81 has been identified as the cytoplasmic proton acceptor^13,18^, although this same residue was found to be non-essential in the bacterial AntR^14^. Both paSzR and psSzR contain a conserved glutamate at the equivalent position (Glu75 in paSzR and Glu76 in psSzR, Supplementary Fig. S1), suggesting this residue may similarly function as the cytoplasmic proton acceptor in these variants. However, the non-essential nature of this glutamate in AntR indicates that alternative or redundant proton acceptance pathways may exist in bacterial SzRs. The extracellular proton donor pathway that reprotonates the RSB also remains to be fully characterized. The conserved cysteine residue in TM3^13,14,18^ (Cys69 in paSzR and Cys70 in psSzR) occupies a position homologous to the DC gate in channelrhodopsins and similar positions in rhodopsin-cyclases and phosphodiesterases. While the functional role of this cysteine in SzRs remains to be fully elucidated, its conservation across the family suggests involvement in ion selectivity or gating mechanisms.

The photocycle characteristics of psSzR and psSzR reveal conserved features and specific variations. Both proteins exhibit the canonical K, L, and M photointermediates characteristic of microbial rhodopsins. However, the overall photocycle turnover is notably slower than SzR1 and SzR4 from *Promethearchaeia* (“*Ca*. Lokiarchaeia”), potentially explaining the reduced H^+^ transport rates observed in whole-cells assay. Notably, the accumulation of the M intermediate is considerably smaller compared to other characterized SzRs^13,14,16–18^, indicating reduced deprotonation of the RSB, which is consistent with the lower H^+^-pumping activity. The substantially faster photocycle of paSzR, combined with its predominantly all-*trans* retinal configuration even under illumination, represents distinct kinetic properties between these variants. Whether these differences reflect adaptations to specific environmental conditions such as light availability or temperature in their native habitats remains to be determined.

Our systematic mutagenesis of conserved residues in TM3 and TM6 clarifies the structural determinants of spectral tuning and proton transport efficiency in bacterial SzRs. The different responses of paSzR and psSzR to equivalent mutations highlights the divergence between these related proteins. The TM3 mutations (paSzR P70S and psSzR A71S) produce reductions in H^+^ transport activity without abolishing function, suggesting that these positions contribute to optimal proton transfer kinetics but are not essential. Structural comparison reveals distinct molecular environments at these positions across SzR subfamilies. In SzR4, Ser76 is positioned near the cytoplasmic acceptor Glu81 and is surrounded by hydrophilic residues including Asn100 and Ser101 in TM4 (Fig. 5, cyan representation). In contrast, Pro70 in paSzR occupies a more hydrophobic environment, flanked by Leu71, Leu72, and Ile73 in TM3 (Fig. 5, magenta representation). This distinct local architecture likely explains why the P70S mutation significantly impacts paSzR activity: the introduction of polar serine into this predominantly hydrophobic microenvironment may disrupt the precise packing and helical geometry required for efficient proton transfer. The proline at position 70 in paSzR confers greater helical rigidity than alanine in psSzR (position 71, Fig. 5, green representation) or serine in SzR4 (position 76, Fig. 5, cyan representation), and its replacement with the more flexible serine may alter the TM3–TM6 interface dynamics without completely disrupting the proton transfer pathway to the conserved glutamate acceptors (Glu75/Glu76 in paSzR/psSzR). TM6 glycine positions (Gly148 in paSzR, and Gly149 in psSzR) emerge as critical determinations of both spectral properties and functional integrity, though with strikingly different consequences between subfamilies. The dramatic phenotype of paSzR G148F, characterized by a 6 nm blue shift and severely impaired proton pumping, demonstrates that this position is critical for maintaining proper retinal geometry. Structural analysis reveals that Gly148/Gly149 in paSzR/psSzR are positioned in close proximity to the retinal β-ionone ring, surrounded by Trp149/Trp150, Phe181/Phe182, Tyr144/Tyr145, and Thr94/Thr95 (Fig. 5, magenta and green representations, respectively). Structural comparison reveals a key difference: SzR4 contains Ile 186 in TM7, whereas paSzR and psSzR have a bulkier phenylalanine residues at the corresponding positions (Phe181 and Phe182, respectively; see Supplementary Fig. S1 and Fig. 5). The introduction of phenylalanine at the normally flexible glycine position (G148F in psSzR and G149F in psSzR) creates severe steric clashes with both the retinal and these neighboring aromatic residues (Phe181/Phe182), constraining the helix movements and conformational changes required during the photocycle. This spatial constraint disrupts the precise molecular choreography necessary for vectorial proton translocation, explaining the reduced activity of these mutants and their inability to adopt SzR4-like configuration with Phe153 (Supplementary Fig. S1). Future structural, spectroscopic, and ecological investigations will further illuminate the remarkable diversity of light-energy conversion strategies employed by microbial life.

## Acknowledgments

M.d.C.M. is grateful to the Azrieli Foundation for the award of the Azrieli Fellowship (cohort 2022-2023) and its support during a 3-months research stay at The Institute for Solid State Physics (ISSP, University of Tokyo, Japan). M.d.C.M. also thanks the ISSP for its collaboration and support during this period. This work was supported by grant M.1.B.A TA_000618_UJA23 (to M.d.C.M) funded by Consejería de Universidad, Investigación e Innovación and by ERDF Andalusia Program 2021-2027, JSPS KAKENHI Grants-in-Aid (grants JP23K21092, JP23K18090, JP23H04404, JP24H02268 to K.I. and JP23K05007 to M.K.), JST CREST (grants JPMJCR22N2 to K.I.), and MEXT Promotion of Development of a Joint Usage/ Research System Project: Coalition of Universities for Research Excellence Program (CURE) (grant JPMXP1323015482 to K.I.), the Israel Science Foundation (Research Center grant 3131/20 to O.B.), and the Nancy and Stephen Grand Technion Energy Program (GTEP). O.B. holds the Louis and Lyra Richmond Chair in Life Sciences. Technical and human support provided by Centro de Instrumentación Científico-Técnica (CICT)-Servicios Centrales de Apoyo a la Investigación (SCAI)-Universidad de Jaén (UJA, MICINN, Junta de Andalucía, FEDER) is gratefully acknowledged.

## Author contributions

M.d.C.M., O.B. and K.I. conceived the project. M.d.C.M. performed molecular biology, protein expression in *E. coli*, ion-pump assay measurement, measurement of *λ*^amax^, protein purification, HPLC analysis of retinal isomers, pH titration, and experimental data analysis. M.d.C.M. and K.I. performed laser-flash photolysis. M.d.C.M. constructed psSzR and paSzR mutants. K.M. performed ion-pump assay measurements and measurement of *λ*^amax^ of psSzR and paSzR mutants. A.R. performed bioinformatics. M.d.C.M., A.R., and K.I. wrote the paper with input from all authors.

