## Supplementary Material for "The bacterial schizorhodopsins: novel light-driven inward proton pumps from Antarctic *Minisyncoccota* (Patescibacteria) and cyanobacteria, with implications for the proton-pumping mechanism"

**Supplementary File S1. Curated and collected SzR sequences.** Spreadsheet containing metadata for the previously expressed and novel sequences, including sequences, taxonomic identity and cluster assignment. Excel spreadsheet.

**Supplementary File S2. Phylogenetic tree of the SzR family.** Phylogenetic tree of the 99% identity cluster representatives: SzRs and outgroups. Numbers next assigned to the nodes indicate support values: Shimodaira-Hasegawa-like approximate likelihood ratio test and Ultrafast bootstrap separated by slash. Newick format.

**Amino acid sequence of paSzR and psSzR**

**>paSzR**
**ORF:Ga0105045_10012953_2**
**scaffold:Ga0105045_10012953_2 genome:3300007517_100 OTU:OTU-2547**
MSLILTLGIVLFSLSSLYFLIKPKAGLNSPFLVSITTLVSYVLMLEGGFLVGSPDTGLYW
TRWAFYGLSCPLLIYEISKQLGLDNKQNFTNIFLTAMVMLTGVLSSITSGDYKLAFFAI
SCVIFCKVLYSVFTSKSDQLVRIAPYMILGWSAFPVVFLLSFEGYGLIQNVLAASIYLG
LDLFTKILFYFHHSAVSKTVTD

**>psSzR**
**ORF:Ga0136636_10000316_21**
**scaffold:Ga0136636_10000316 genome:3300012044_11 OTU:OTU-4691**
MNIVLLAGIVMFACSSLYFWLTSKKEFNSAFLVSFITLISYIIMYEGRLLIGDSETGLYW
TRWAFYGLSCALLTYEISKQLNIAKGIQWLAIILTPIVMFTGSLASIYTETYKWIWFVIS
SIAFLAIAKIYYSTKSAELPRISMYFLFGWSVFPLVFLLSPEGLSIINNSFAGTVYLVLD
FFTKIIFYIQYSNLHKKL

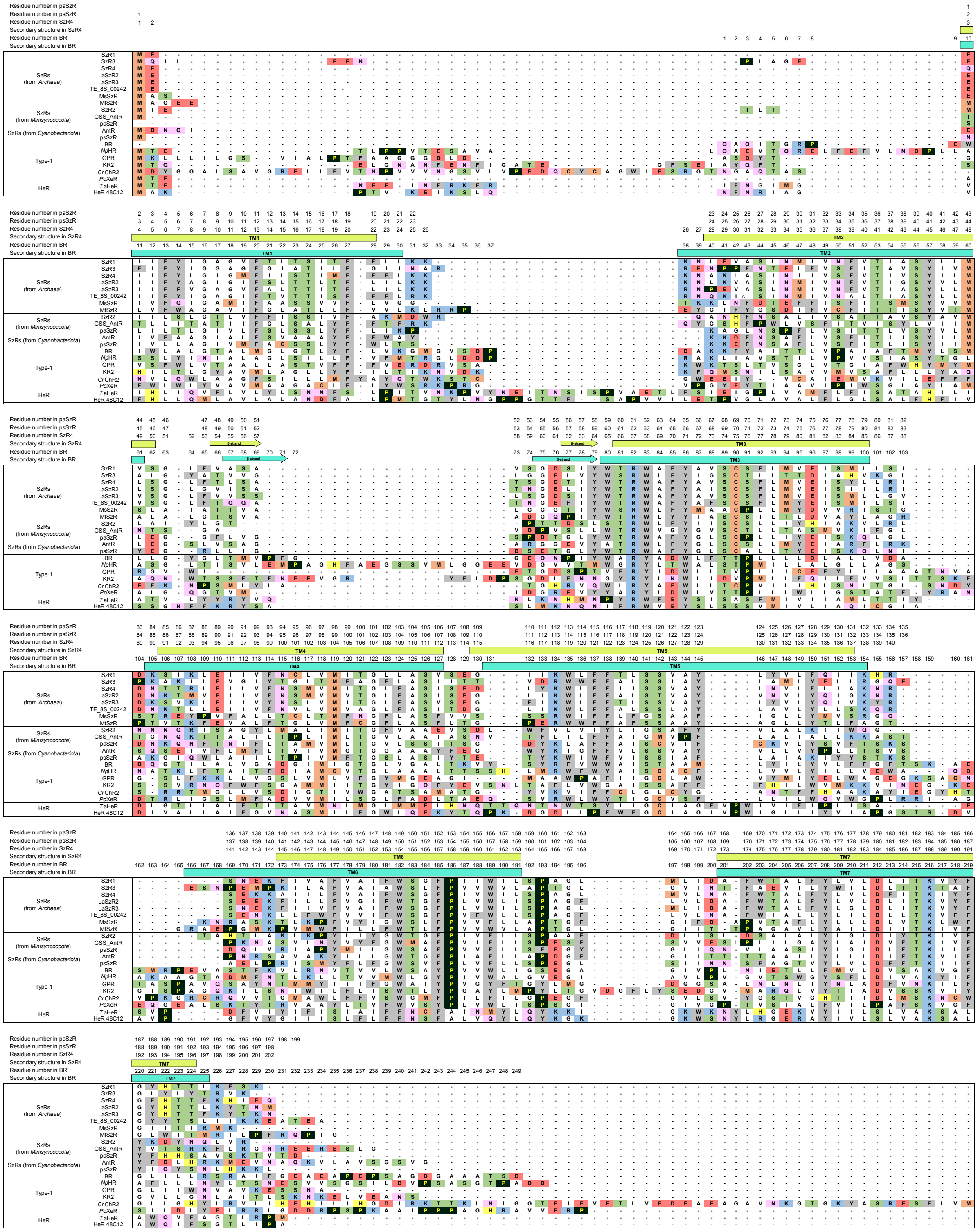


**Supplementary Fig. S1. Multiple amino acid sequence alignments**. The amino acid sequences were aligned using PROMALS3D[1] and MEGA11[2]. SzR1, SzR3, SzR4, LaSzR2, LaSzR3, TE_8S_00242, MsSzR, MtSzR, SzR2, Antarctic rhodopsins (GSS_AntR and AntR), bacteriorhodopsin (BR), *Natromonas pharaonis* halorhodopsin (*Np*HR), green-absorbing proteorhodopsin (GPR), *Krokinobacter* rhodopsin 2 (KR2), *Chlamydomonas reinhardtii* channelrhodopin 2 (*Cr*ChR2), *Parvularcula oceani* xenorhodopsin (*Po*XeR), and HeRs (*Thermoplasmatales* archaeon SG8-52-1 heliorhodopsin (*Ta*HeR) and HeR48C12) were aligned with the sequences of paSzR and psSzR. The residue numbers of paSzR, psSzR, SzR4, and BR are shown on top of the residues, and the positions of TM helices and beta-strands, based on X-ray crystallographic structures of SzR4 (PDB ID: 7E4G[3]) and BR (PDB ID: 7Z09[4]), are indicated by rectangles and arrows (green for SzR4 and blue for BR), respectively. For clarity, C-termini and interhelical loops of BR, *Np*HR, GPR, KR2, *Cr*ChR2, *Po*XeR, *Ta*HeR, and HeR 48C12 were omitted.

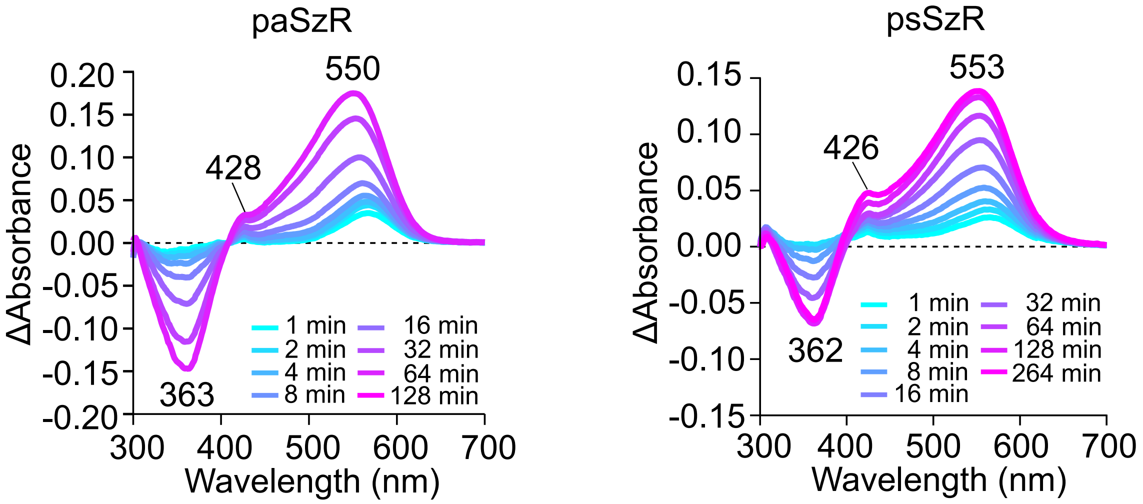


**Supplementary Fig. S2. Light-induced difference absorption spectra of paSzR and psSzR rhodopsins.** Difference absorption spectra before and after HA bleaching reactions of paSzR and psSzR rhodopsins in solubilized *E. coli* membranes. The $\lambda^{a}_{max}$ values were determined by the positions of the absorption indicated in each panel and the absorption of retinal oxime produced by the hydrolysis reaction of RSB and HA observed as peaks in the proximity of 360–370 nm at pH 7.0. The samples were exposed to light for up to 128–264 min.

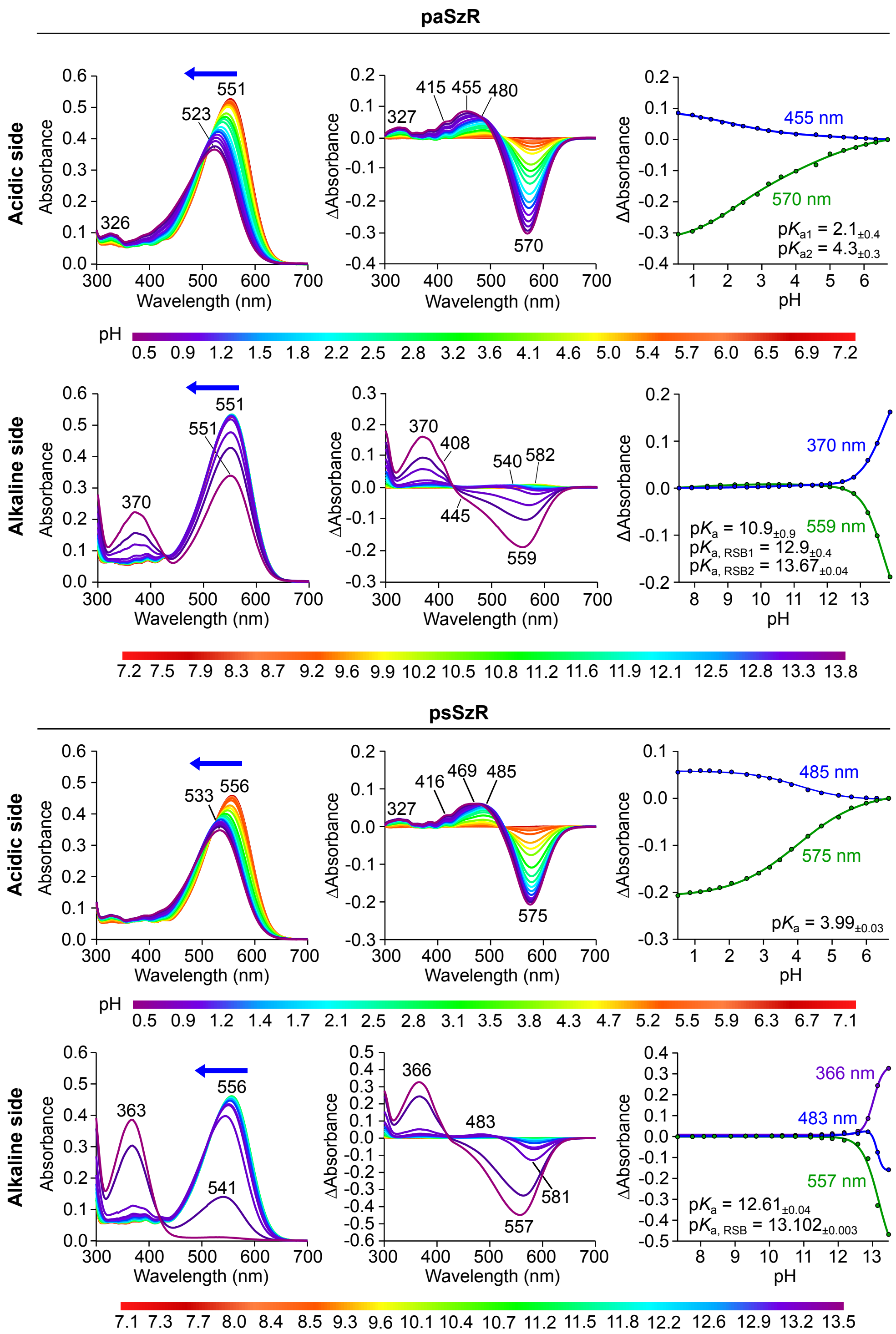


**Supplementary Fig. S3. pH dependence on the absorption**. Absorption spectra (left), different absorption spectra (center), and pH titration curves for the calculation of p$K_a$ (right) were measured depending on acidic and alkaline pH changes. The blue-shifted (blue arrow) absorption spectra at each pH are indicated by arrows. The pH titration curves were analyzed using the Henderson–Hasselbalch equation[5]. While denaturation of the paSzR and psSzR proteins would result in the exposure of its RSB chromophore to the environment and the RSB deprotonation under the extreme pH conditions, all spectra retained visible-absorbing components (except at very alkaline pH (pH > 13.5) for psXR), indicating that paSzR and psSzR maintained its native protein folding over the entire pH range examined in this study.

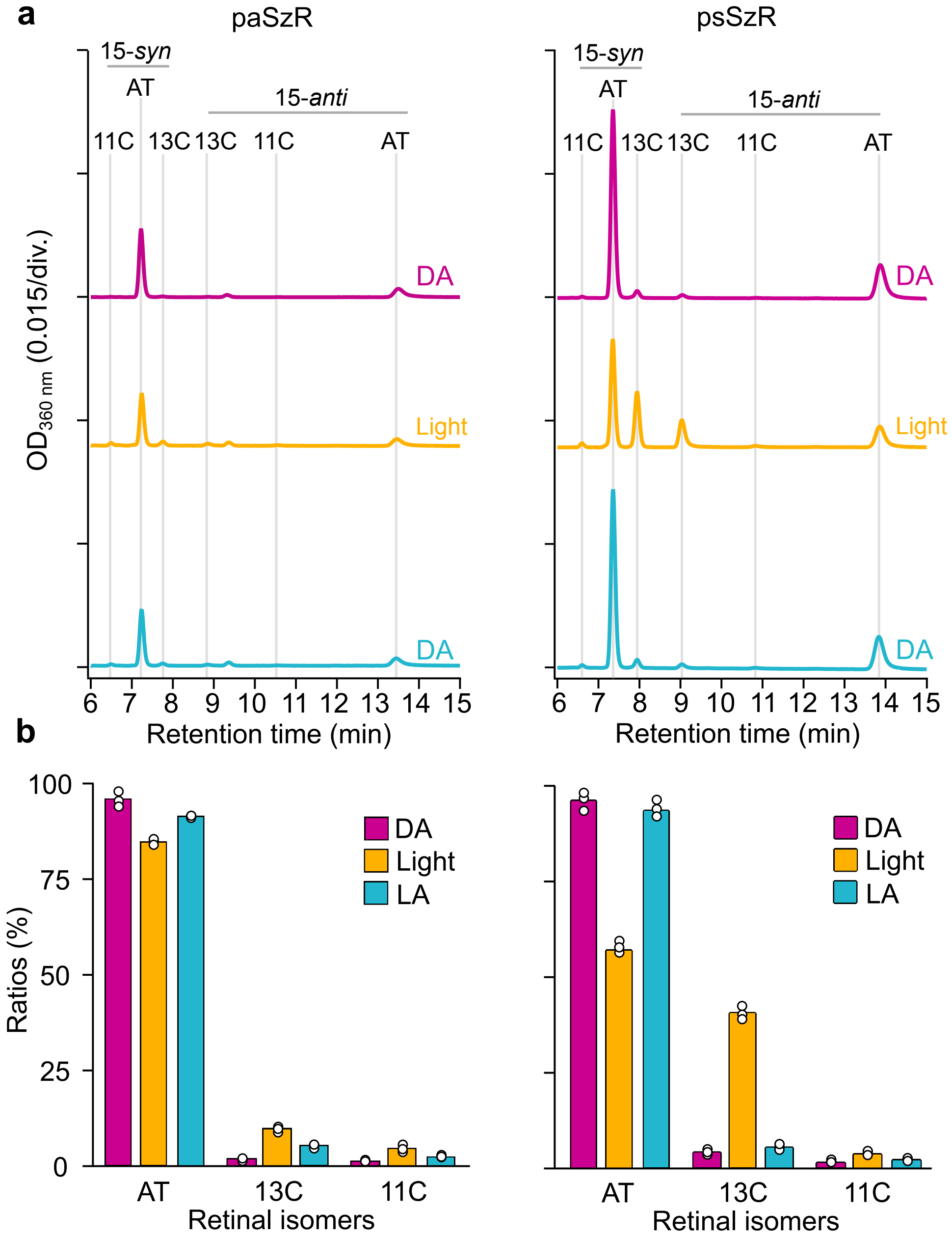


**Supplementary Fig. S4. HPLC analysis of the retinal configuration**. **a** Chromatogram of HPLC analyses and **b** the compositions of the retinal isomers under the dark (DA, violet), light (orange), and light-adapted (LA, blue) conditions, where AT, 13C, 11C, *syn*, and *anti* indicate all-*trans*, 13-*cis*, 11-*cis*, *syn*, and *anti* configurations, respectively. The corresponding data are listed in Supplementary Table S2.

**Supplementary Table S1. List of primers used for site-directed mutagenesis.**

| Mutations | Sense primer | Antisense primer |
|---|---|---|
| paSzR P70S | tttacggtctgtcctgctcgctgctgatttatgaa | ttcataaatcagcagcgagcaggacagaccgtaaa |
| paSzR G148F | gctccgtatatgattctgttttggtcggcgtttccggt | accggaaacgccgaccaaaacagaatcatatacggagc |
| psSzR A71S | tttctacggtctgtcttgcagtctgctgacctacgaaatc | gatttcgtaggtcagcagactgcaagacagaccgtagaaa |
| psSzR G149F | ctctatgtacttcctgttcttttggtctgttttcccgctg | cagcgggaaaacagaccaaaagaacaggaagtacatagag |

**Supplementary Table S2. Retinal configuration in psSzR and paSzR.** The composition of the retinal isomers was determined through HPLC analysis for retinal oxime produced by the hydrolysis of the retinylidene Schiff base with hydroxylamine. AT, 13C, and 11C indicate all-*trans*, 13-*cis*, and 11-*cis* configurations, respectively.

| Protein | Light conditions | AT (%) | 13C (%) | 11C (%) |
|---|---|---|---|---|
| paSzR | DA | 96 ± 1 | 2.1 ± 0.8 | 1.7 ± 0.2 |
| | Light | 84.0 ± 0.2 | 9.8 ± 0.1 | 5.17 ± 0.09 |
| | LA | 91.62 ± 0.01 | 5.3 ± 0.2 | 3.1 ± 0.2 |
| psSzR | DA | 95.6 ± 0.3 | 3.6 ± 0.2 | 0.73 ± 0.02 |
| | Light | 56.9 ± 0.4 | 40.3 ± 0.5 | 2.7 ± 0.2 |
| | LA | 93.5 ± 0.2 | 4.89 ± 0.06 | 1.6 ± 0.1 |
